# UbiFast, a rapid and deep-scale ubiquitylation profiling approach for biology and translational research

**DOI:** 10.1101/785378

**Authors:** Namrata D. Udeshi, Deepak C. Mani, Shankha Satpathy, Shaunt Fereshetian, Jessica A. Gasser, Tanya Svinkina, Benjamin L. Ebert, Philipp Mertins, Steven A. Carr

## Abstract

Protein ubiquitylation is involved in a plethora of cellular processes. Defects in the ubiquitin system are at the root of many acquired and hereditary diseases. While antibodies directed at ubiquitin remnants (K-ε-GG) have improved the ability to monitor ubiquitylation using mass spectrometry, methods for highly-multiplexed measurement of ubiquitylation in tissues and primary cells using sub-milligram amounts of sample remains a challenge. Here we present a highly-sensitive, rapid and multiplexed protocol for quantifying ∼10,000 ubiquitylation sites from as little as 500 ug peptide per sample from cells or tissue in a TMT10 plex in ca. 5 hr. High-field Asymmetric Ion Mobility Spectrometry (FAIMS) is used to improve quantitative accuracy for posttranslational modification analysis. We use the approach to rediscover substrates of the E3 ligase targeting drug lenalidomide and to identify proteins modulated by ubiquitylation in models of basal and luminal human breast cancer. The sensitivity and speed of the UbiFast method makes it suitable for large-scale studies in primary tissue samples.

## Introduction

Protein ubiquitylation is an important and highly conserved post-translational modifications (PTM) essential for regulating protein-turnover through the ubiquitin-proteasome system. Ubiquitylation occurs most commonly on the ε-amino group of protein lysine residues through the concerted action of activating (E1), conjugating (E2), and ligating (E3) enzymes^1,2^. The E3 ligases attach ubiquitin chains to specific substrate proteins, often targeting these proteins for degradation via the proteasome. Mutations and other changes that result in dysregulation of either ligases or deubiquitinases (DUBs) may lead to aberrant activation or deactivation of pathways involved in many disease processes, notably cancer progression and metastasis, immune disorders and neurological diseases among others. Their role in oncogenesis has been well-described^3-7^. The potential druggability of E3 ligases with small molecules such as lenalidomide as treatments for a variety of cancers has greatly increased interest by the biotechnology and pharmaceutical industries in this class of enzymes^5,8^.

Historically, identification of protein ubiquitylation sites by mass spectrometry has proven to be challenging because of the low stoichiometry of ubiquitylated proteins, the size of the modification itself, and the diversity in resulting ubiquitin chain types. Robust, large-scale detection of endogenous ubiquitylation sites by mass spectrometry requires a technique that facilitates specific enrichment of only the modified lysine containing peptides of ubiquitylated substrate proteins. To this end, global analysis of protein ubiquitylation has markedly improved with the commercialization of antibodies specific for the di-glycyl remnant produced on ubiquitylated lysine residues (K-ε-GG) following trypsin digestion^9-12^. Specifically, trypsin digestion of ubiquitylated proteins cleaves off all but the two C-terminal glycine residues of ubiquitin from the modified protein. These two C-terminal glycine residues remain linked to the epsilon amino group of the modified lysine residue in the tryptic peptide derived from digestion of the substrate protein (**Figure 1A**). The presence of the di-glycine group on the sidechain of lysine prevents cleavage by trypsin at that site, resulting in an internal modified lysine residue in a formerly ubiquitylated peptide. This K-ε-GG group is recognized and enriched using an anti-K-ε-GG antibody (**Figure 1A**).

**Figure 1.**
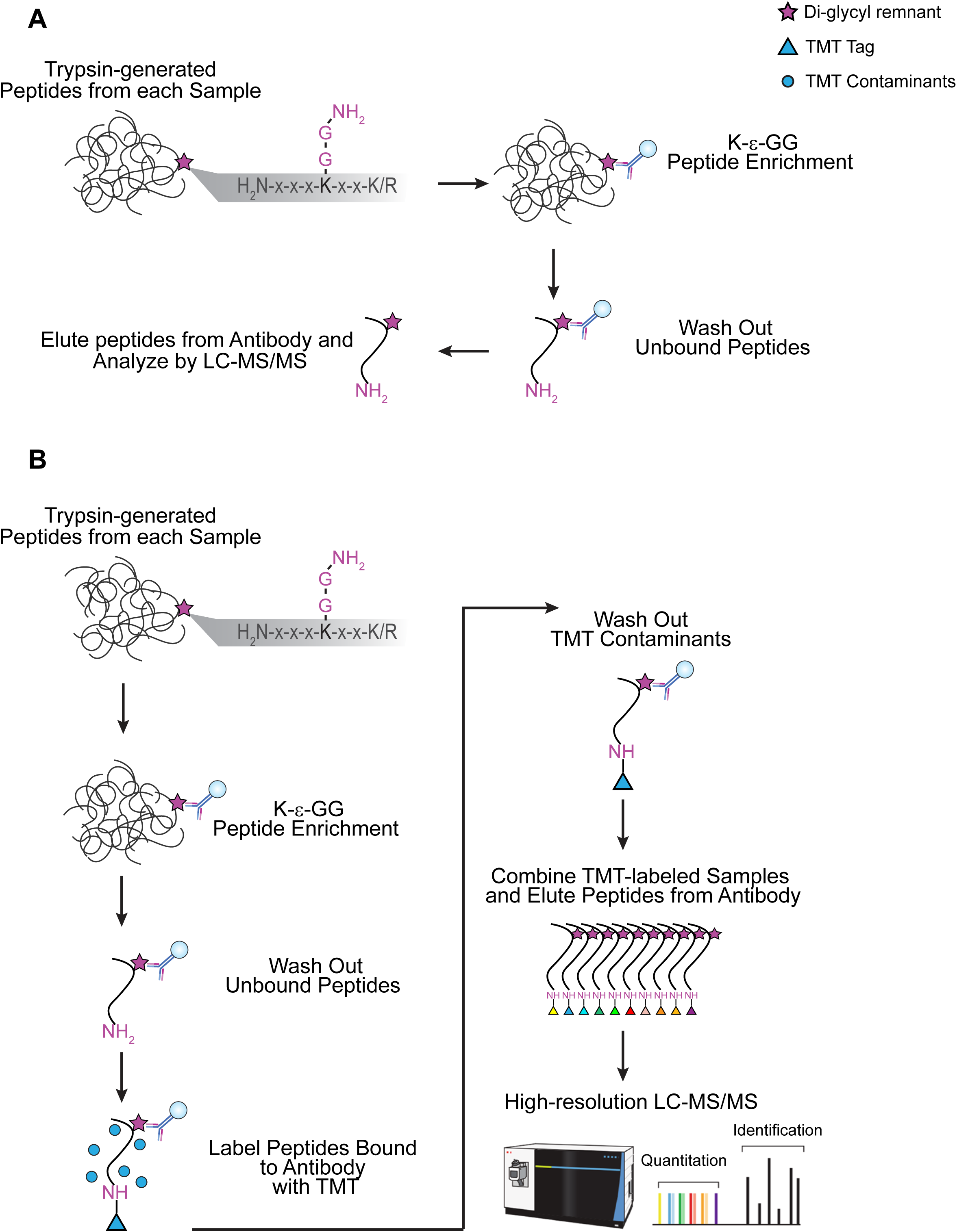
Comparison of methods for multiplexed ubiquitylome analysis A) Conventional workflow for an anti-K-ε-GG antibody-based enrichment without isobaric labeling, where the antibody recognizes primary amine containing K-ε-GG epitope. B) UbiFast workflow employing on-antibody labeling with isobaric mass tag reagents (e.g., TMT10/11) for anti-K-ε-GG antibody-based enrichment of ubiquityxslated peptides.

To enable precise relative quantification of ubiquitylated peptides and sites across differing samples under perturbation conditions, SILAC (Stable Isotope Labeling by Amino Acids in Cell Culture) has been historically used to differentially label cells grown in culture prior to antibody enrichment. SILAC enables comparison of ubiquitylation sites from up to three samples in a single experiment. We and others have successfully applied this approach in a range of quantitative biological studies using metabolically-labeled cells ^5,13-18^.

Isobaric chemical tags, such as the Tandem Mass Tag (TMT) system, offer many advantages over SILAC for quantifying post-translational modifications by mass spectrometry. They facilitate a decrease in starting material used for enrichment, enable comparison of 11 or more conditions in a single experiment, and significantly minimize the number of missing peptides detected and quantified across all experimental conditions relative to label-free or SILAC-based experiments. However, a major limitation of the ubiquitylation profiling approach described above has been that the commercially available antibodies used to recognize and enrich peptides having the di-glycyl remnant on the side-chain of lysine do not work when the N-terminus of the di-glycyl remnant is derivatized with either iTRAQ or TMT. Efforts to produce antibodies that selectively enrich either an iTRAQ- or TMT-derivatized K-ε-GG peptide and not other iTRAQ or TMT-labeled peptides have failed to date (unpublished). Until recently this limitation has restricted use of ubiquitin profiling to fast growing cell lines that can be metabolically labeled in culture and has prevented multiplexed, quantitative analysis of human or animal-derived tissues or primary cell culture models using isobaric reagents.

To address the need for profiling ubiquitylation sites in tissue samples, Rose et al. introduced a method where samples are enriched at the peptide level using the anti-K-ε-GG antibody prior to labeling with TMT10 reagents^19^. After elution from the antibody and labeling, the enriched peptides are subjected to high pH reversed-phase chromatography using spin columns, fractionated into six fractions, and analyzed by LC-SPS-MS3. In this study, 5,000 - 9,000 ubiquitylated peptides were quantified in cells using 1 mg sample/TMT label state and in tissue using 7 mg of sample/TMT label state using 18 hr of instrument time. This approach represented a breakthrough in the ability to employ isobaric chemical tagging for ubiquitylation profiling of tissues. However, the large amount of sample and lengthy analysis time required to achieve significant depth of detection and quantification of the ubiquitylome in tissue using this approach would likely preclude its use for analysis of more limited samples such as primary cells and human tumor samples.

The limitations noted above led us to consider other possible approaches to overcome the inability to use anti-K-ε-GG antibodies with TMT labeling. Mass spectrometry (MS) has previously been used to map the linear amino acid epitope of a protein recognized by an antibody (Ab)^20,21^. In these studies, protein in free and antibody-bound forms was proteolytically digested and the resulting peptides analyzed by MS. The epitope was protected from proteolysis when the antibody was bound and could be identified on the basis of the differential MS analysis. Inspired by this earlier work, we hypothesized that the di-glycyl remnant of ubiquitylated peptides would not be solvent exposed when bound to the anti-K-ε-GG antibody. Leveraging this approach, we have developed a method that increases both sensitivity and throughput for highly-multiplexed ubiquitylation profiling. In this approach we term UbiFast, K-ε-GG peptides are labeled with TMT reagents while still bound to the anti-K-ε-GG antibody. By doing so, the amine-reactive, NHS-ester group of the TMT reagent reacts with peptide N-terminal amine group and the ε-amine groups of lysine residues, but not the primary amine of the di-glycyl remnant. TMT-labeled K-ε-GG peptides from each sample are combined, eluted from the antibody, and analyzed by single-shot, high-performance LC-MS/MS **(Figure 1B**). We reasoned that TMT labeling of K-ε-GG peptides bound to the antibody beads could potentially increase sensitivity, reduce the levels of TMT-based contaminant side-products, and avoid the need to offline fractionate prior to MS measurement. We have used this method to profile patient-derived breast cancer xenograft tissue samples in a TMT 10-plex experiment using 0.5 mg input/sample to quantify >10,000 distinct ubiquitylation sites. The method paves the way for quantification of ubiquitin remnants in human tissues and more physiological patient-derived cell models where sample amounts are limiting.

## Results

### Optimization of On-antibody TMT Labeling and Comparison of On-antibody TMT labeling with In-solution TMT labeling for Ubiquitylome Analysis

To establish the feasibility of labeling peptides with TMT reagents while bound to the anti-K-ε-GG-antibody and to determine the optimal amount of labeling reagent and labeling time, K-ε-GG peptides were enriched in triplicate from 1 mg of Jurkat peptides cells and labeled with varying amounts of a single TMT reagent for varying durations while peptides were still bound to antibody (**Figure 1B, Supplemental Figure 1, Supplemental Tables 1,2**). We found that 10 min labeling with 0.4 mg of TMT reagent provided the best balance of numbers of identified TMT-labeled K-ε-GG peptides and completeness of labeling (>92%) for K-ε-GG peptides bound to anti-K-ε-GG antibody. In order to prevent potential TMT cross-labeling when samples are combined, it is important that the labeling reaction be fully quenched. Therefore, we tested quenching of the TMT reactions with 5% hydroxylamine. The quenching was successful in stopping the labeling reaction, evidenced by a slightly reduced labeling efficiency. Importantly, quenching also increased the number of K-ε-GG peptides identified by almost 10% (**Supplemental Figure 2, Supplemental Table 3A**). We find that label free analysis of enriched K-ε-GG peptides results in lower relative yield of K-ε-GG peptides when compared to K-ε-GG enrichment coupled to TMT labeling (**Supplementary Figure 2, Supplemental Table 3B).**

We next compared in-solution TMT labeling of K-ε-GG peptides to on-antibody TMT labeling of K-ε-GG peptides with respect to the total numbers of K-ε-GG peptides detected, the relative yield of K-ε-GG peptides (relative yield is the percentage of K-ε-GG peptides relative to the total peptides identified in the sample) and the efficiency of TMT labeling (**Supplemental Figure 3 A-D**, respectively). Jurkat peptide samples (1 mg, each) were enriched with the anti-K-ε-GG antibody and peptides were labeled with a single TMT reagent, either while the peptides were bound to the antibody or using the in-solution labeling method previously described^19^ (**Supplemental Figure 3A**). On-antibody TMT labeled samples resulted in 6087 K-ε-GG PSMs with a relative yield of 85.7%, while samples labeled in-solution resulted in 1255 K-ε-GG PSMs with a relative yield of 44.2% (**Supplemental Figure 3B, C, Supplemental Table 4**). The labeling efficiency of on-antibody and in-solution TMT labeling methods were both high, with 98% of peptides being at least partially labeled (**Supplemental Figure 3D**).

As both labeling methods are designed to be used with multiplexed samples, we carried out a head-to-head comparison where both labeling methods were used to analyze 10 process replicates of peptides from HeLa cells, with 1 mg peptide input per replicate (**Supplemental Figure 4**) using single-shot LC-MS/MS methods with 2 injections per sample to and longer gradients than in the initial experiments described above (154 min/injection vs. 110 min/injection) totaling to 5.1 h of instrument time. We find that injecting samples twice leads to a moderate boost in identifications. In concordance with the single sample experiments, the on-antibody labeling method identified many more fully quantified, distinct K-ε-GG peptides compared to the in-solution labeling method (9069 vs. 4587 peptides) (**Supplemental Figure 4B and Supplemental Tables 5A,B**). The relative yield of K-ε-GG peptides was 85.4% using the on-antibody labeling approach compared to 49.9% for the in-solution labeling method (**Supplemental Figure 4B**). There was significant overlap in the commonly identified sites, with approximately 80% of the sites identified using the in-solution labeling method also identified by the on-antibody labeling approach (**Supplemental Figure 4C**). The median CVs between the 10 process replicates were very similar for both TMT labeling methods (**Supplemental Figure 4D**). TMT labeling of K-ε-GG peptides bound to the antibody beads could also potentially reduce the levels of TMT-based contaminant side-products in the final sample, which often show up as +1 precursors ^22^, and avoid the need to offline fractionate prior to MS measurement. Singly-charged precursors constituted a far lower percent of the total precursor ion current using on-antibody TMT labeling than in-solution labeling (3.9% vs. 16.5% (**Supplemental Figure 4E, Supplemental Table 5C,5D**). These results demonstrate that on-antibody TMT labeling more effectively removes TMT contaminants and improves K-ε-GG enrichment specificity which is what likely leads to an increase in identified and fully quantified K-ε-GG peptides.

### Identification of Lenalidomide Targets in Multiple Myeloma Cells

In prior work, we employed ubiquitylation profiling by mass spectrometry in SILAC-labeled multiple myeloma cell lines to reveal the mechanism of action of lenalidomide, an anti-tumor drug in multiple myeloma^5^. This study, which utilized 10 mg of peptide/SILAC state, revealed that lenalidomide causes degradation of the Ikaros transcription factors IKZF1 and IKZF3. To benchmark our new approach, we repeated our previously published SILAC-based experiment by treating MM1S cells with 1 uM lenalidomide for 12h and MG-132 for 3h, or with only MG-132 for 3h using either HCD-MS2, SPS-MS3^22,23^ or FAIMS-MS2, a method recently shown to increase the numbers of detected and quantified peptides in proteome studies^24,25^ (**Figure 3**). We then enriched K-ε-GG peptides and carried out on-antibody TMT10 labeling for ubiquitylome profiling as described above using just 1 mg of peptide input per sample. The final enriched TMT10-plex sample mixture was divided into thirds and one-third of the sample was analyzed by each of the three data acquisition methods (**Figure 2A**). Using the MS2 approach we quantified 15,612 distinct ubiquitylation sites (**Figure 2B, 2C and Supplemental Table 6A**). Using FAIMS-MS2, the number of quantified ubiquitylated peptides was similar (15,166) (**Supplemental Table 6A**). In contrast, only 3970 ubiquitylated peptides were observed using SPS-MS3 (**Supplemental Table 6A**), more than three-fold fewer than using either MS2 or FAIMS-MS2. The reproducibility of all three methods was high, with median coefficients of variation across process replicates of <20% (MS2: 13.2%, FAIMS-MS2:15.8%, SPS-MS3: 10%).

**Figure 2.**
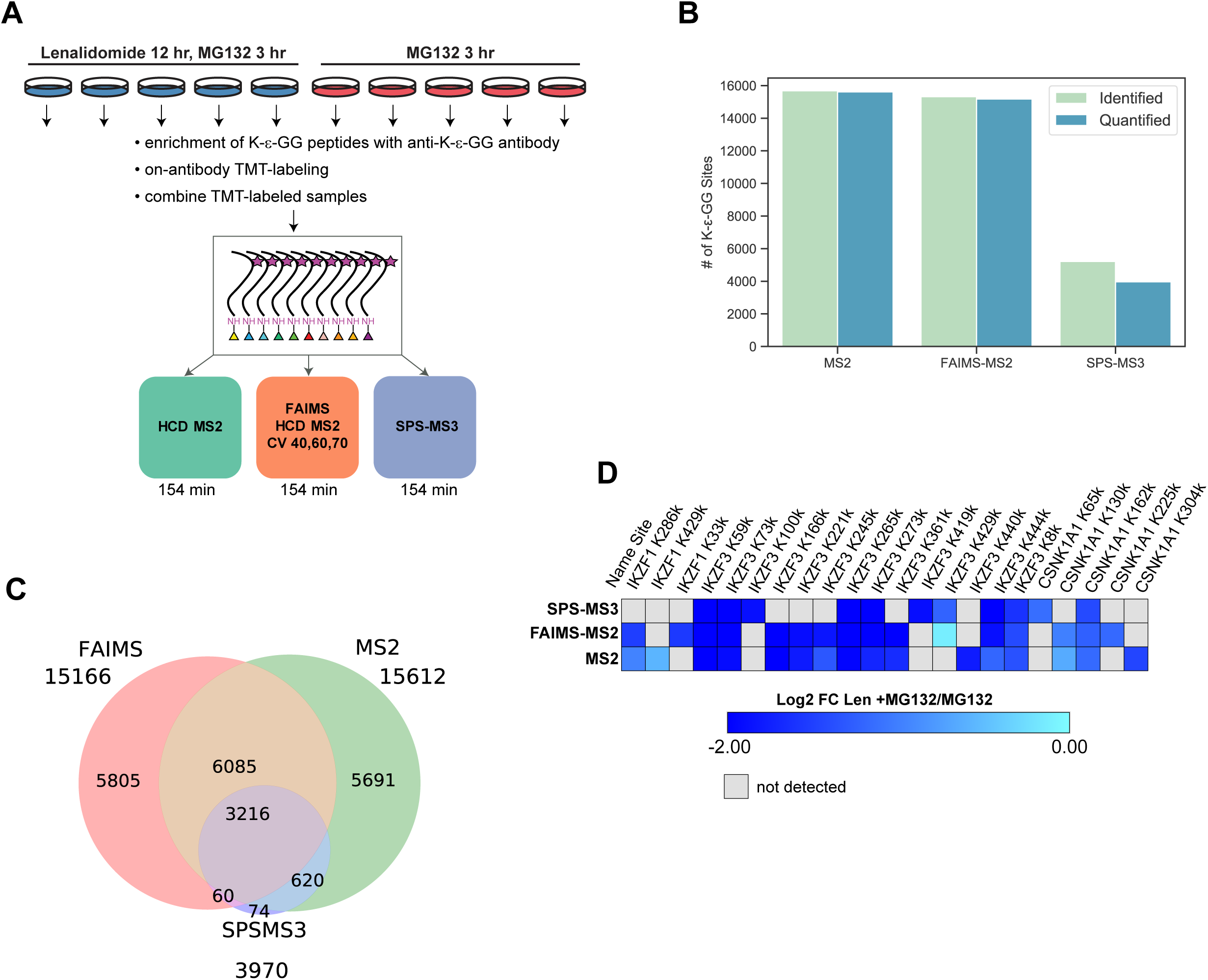
Benchmarking UbiFast on-antibody K-ε-GG enrichment and TMT labeling in lenalidomide-treated MM1S cells using HCD-MS2, SPS-MS3 and FAIMS-MS2. A) Experimental design for TMT10-based assessment of global changes in ubiquitylation and protein levels B) Bar plots show the number of distinct quantified K-ε-GG sites identified in each type of MS experiment. C) Venn diagram shows the overlap of quantified K-ε-GG sites across each MS experiment type (HCD-MS2, SPS-MS3, FAIMS-MS2) D) Venn diagram shows the overlap of quantified K-ε-GG sites from IKZF1, IKZF3, and CSNK1A1 across each MS experiment type (HCD-MS2, SPS-MS3, FAIMS-MS2). E) Heatmap of log2 fold change values for 12 h lenalidomide + 3h MG-132 versus 3 h MG-132 treated MM1S cells for IKZF1, IKZF3, and CSNK1A1. HCD-MS2 and FAIMS-MS2 data have been filtered to show only those sites that had >90% precursor isolation purity.

**Figure 3.**
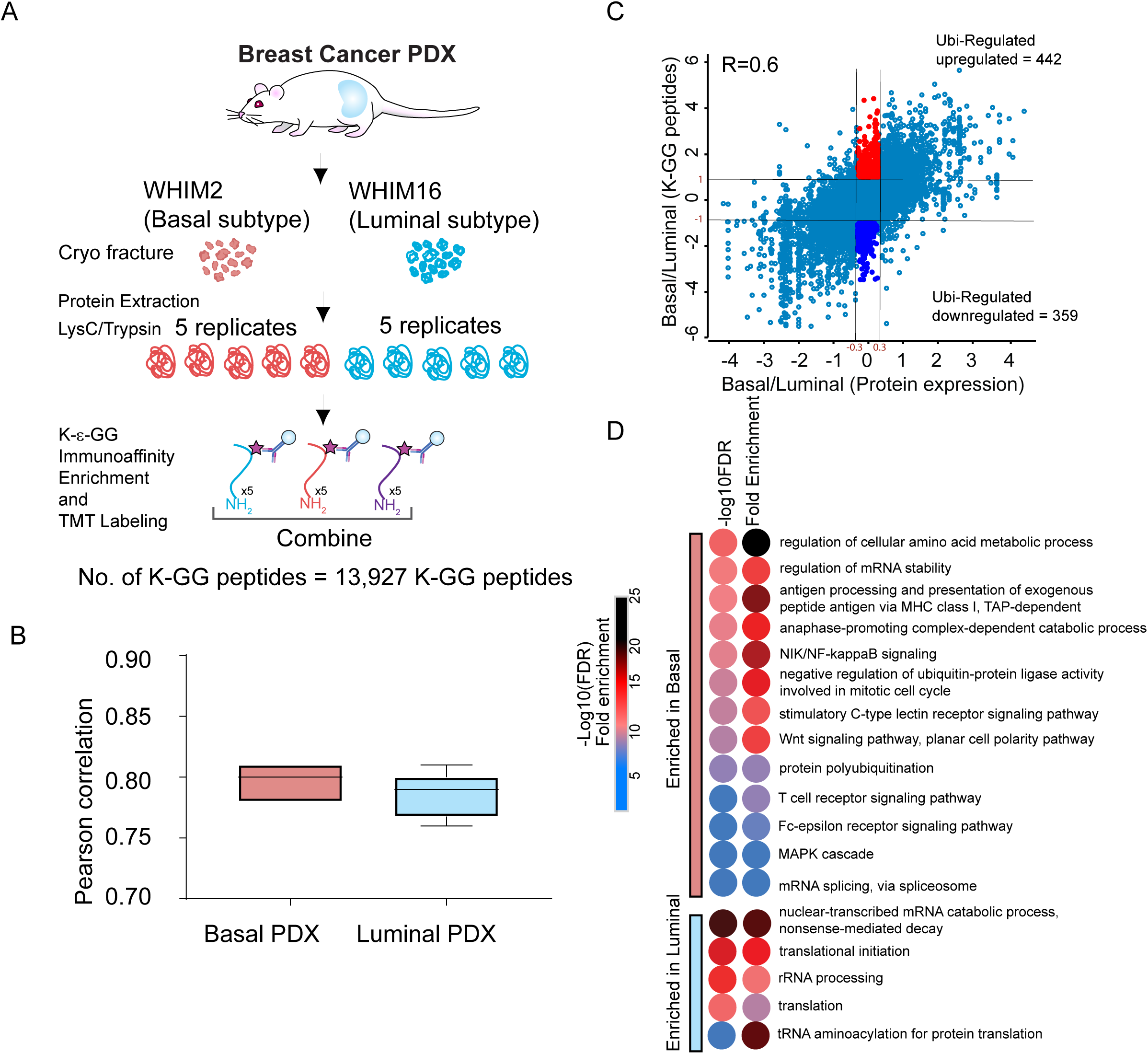
Ubiquitylation analysis of breast cancer PDX models. A) Schematic diagram showing experimental design used to enrich di-Gly modified peptides from the Luminal and Basal breast cancer PDX models. B) Pearson correlation between Luminal and Basal PDX replicates C) Scatter plot showing log2 fold-change between Basal and Luminal PDXs, for expression of ubiquitylated peptides and the corresponding protein. The proteins that were regulated exclusively at the level of their ubiquitylation are color coded. D) Pathway enrichment of proteins regulated primarily at the level of ubiquitylation.

Using either MS2 or FAIMS-MS2, we observed the expected changes in ubiquitylation induced by lenalidomide treatment on multiple lysine residues of IKZF1, IKZF3, and CSNK1A1. Because IKZF1 and IKZF3 are rapidly ubiquitylated and targeted for degradation, the resulting decrease in corresponding ubiquitylated peptides is a result of decreased absolute levels of these proteins ^5^. However, using SPS-MS3, changes in ubiquitylation of one of the key transcription factors, IKZF1, were not detected, and fewer ubiquitylated peptides overall were detected on the IKZF3 and CSNK1A1 (**Figure 2D**). Importantly, the expected ubiquitylated and degraded proteins were identified using only 1 mg peptide/sample compared to a total of 10 mg input/sample used in the SILAC approach^5^. Together, these results support the feasibility and value of multiplexed ubiquitylation profiling in systems where sample amounts are limiting.

### Accuracy of Ubiquitylated Peptide Quantification

Using HCD-MS2 we observed several ubiquitylated peptides from IKZF1, IKZF3, and CSNK1A1 that appeared to not be significantly down-regulated upon lenalidomide treatment despite the observation of other ubiquitylated peptides from these same proteins that were strongly regulated (**Supplemental Figure 5A**). We reasoned that this was most likely the result of ratio compression, an expected and well-described issue related to TMT quantitation^22,26^. Precursor isolation purity filtering is a common approach for removing interferences in TMT experiments^27,28^. Filtering MS2 and FAIMS-MS2 data for only those peptides having > 90% precursor isolation purity (PIP) significantly improved quantification accuracy (**Supplemental Figure 5B**) while decreasing the total number of ubiquitylated peptides observed in MS2 by 40.8% (**Supplemental Table 6B**). As expected, FAIMS-MS2 reduced precursor co-isolation interference and stringent PIP filtering retained relatively more ubiquitylated peptides than stringent PIP filtering of MS2 data, reducing the total number of ubiquitylated peptides by only 28% (**Supplemental Figure 5C and Supplemental Table 6B**). SPS-MS3 had the greatest accuracy, but identified fewer of the statistically significant sites that were seen by either MS2 and FAIMS-MS2 (**Supplemental Figure 5D and Supplemental Table 6**).

### Applying UbiFast for rapid profiling of tumor tissue

To demonstrate the utility of UbiFast approach to quantitatively profile the ubiquitylome of small amounts of tissue, we isolated tumors from two previously described breast cancer patient-derived xenograft (PDX) models, representing basal (WHIM2) and luminal (WHIM16) subtypes of breast cancer, respectively^29^ (**Figure 3A**). These models faithfully reproduce genomic features of the disease, exhibiting distinct expression of key basal and luminal genes, and proteome driven basal and luminal pathway signatures^30^. Ubiquitylated peptides were enriched from 5 replicates each of WHIM2 and WHIM16 using 0.5 mg of peptide per sample. As described above, peptides enriched using anti-K-ε-GG antibody were TMT labeled on antibody. TMT-labeled peptides from each sample were then combined and analyzed using two single-shot LC-MS/MS runs. FAIMS was used with two compensation voltage (CV) sets per injection (see Methods) to reduce ratio compression and increase the total number of ubiquitylated peptides identified. We find that injecting samples twice using a different compensation set for each analysis results in lower peptide overlap (and therefore higher total identifications) relative to two independent injections without FAIMS. A total of 13,501 human K-ε-GG peptides were observed, 81% (10,942) of which were quantified across all 10 TMT channels (**Supplemental Table 7A**). The correlation across replicates was high, with a median Pearson correlation of 0.78 - 0.81 (**Figure 3B**).

To identify the relationship between ubiquitylated peptides and their protein expression, we compared ubiquitylome to a previously published deepscale proteomics dataset obtained on the same basal and luminal PDX breast cancer models that used a similar TMT10 design with analysis of 24 offline basic reverse phase fractions (vs. UbiFast using single-shot methods) using a similar TMT10 workflow^31^ (**Supplemental Table 7C**). There was a high degree of correlation between the fold-changes observed between basal and luminal expression at the level of K-ε-GG modified peptides and the corresponding protein abundance changes. This further supports robust quantification of ubiquitylated peptides relative to deep-scale fractionated proteome as it is expected that the majority of changes observed in the ubiquitylome will be driven by changes in the levels of the corresponding proteins (Pearson correlation = 0.6) (**Figure 3C**). However, in addition, we also identified a total of 801 K-ε-GG peptides that were regulated (log2 fold-change >1 and <-1) primarily at the level of ubiquitylation (referred to as Ubi-regulated), but that did not show regulation at the protein level (log2 fold-change >-0.3 and <0.3) (**Figure 3C, Supplemental Table 7D**). A total of 152 sites showed marked outlier regulation with log2 fold-change >2 and <-2 (**Figure 3C, Supplemental Table 7D**). Of note, we observed differential ubiquitylation of 15 E3 ligases; TRAF7, HUWE1, NEDD4L, UBE3A, UBE4B, CBL, BIRC6, RAD18, RNF146, MIB1, HERC2, AFF4 that showed upregulated ubiquitylation in Basal subtype whereas ITCH, TRIM25, RNF185 and IRF2BPL1 showed upregulated ubiquitylation in Luminal PDX model (**Supplemental Table 7D**). Pathway enrichment of Ubi-regulated proteins (**Figure 3D**) showed enrichment of several immune-signaling pathways in the basal PDX models consistent with the critical role of ubiquitylation in immune signaling^14,32,33^. Interestingly, Basal or triple-negative breast cancer (TNBC) shows high immunological activity and are attractive candidates for immuno-oncology therapies^34^ and therefore, upregulated ubiquitylation of immune-signaling is consistent with the “immune hot” environment within the basal PDX tumor.

To begin to evaluate the crosstalk between two distinct lysine modifications, we compared differential ubiquitylation and acetylation (K-Ac) in these basal and luminal PDX models (**Supplemental Figure 6, Supplemental Table 7C**). Enrichment of K-Ac peptides were performed as previously described with modifications ^35^. TMT-labeled peptides from five replicates each of the basal and luminal PDX models were mixed at an equal ratio, fractionated into 4 fractions and subjected to immuno-enrichment using an anti-acetyl antibody (PTM-SCAN acetyl-kit, Cell Signaling Technologies). A total of 11,929 acetylated peptides were identified with 10,967 of these identified across all channels. Over 2550 sites showed both acetylation and ubiquitylation with pearson correlation of 0.45 (**Supplemental Figure 6 A,B, Supplemental Table 7D**). Interestingly, a small subset of Ubi-regulated sites were also inversely regulated by acetylation suggesting a regulatory crosstalk between ubiquitylation and acetylation (**Supplemental Figure 6 C, D**). In summary, this study highlights the feasibility of UbiFast to identify ubiquitylation driven pathways in tumors as well as cell lines, and that it can also be used to investigate the crosstalk of lysine ubiquitylation and other lysine modifications such as acetylation as exemplified in this study. Future studies in a larger number of PDX or human tumors are needed to elucidate in-vivo functional roles.

## Discussion

A major advantage of TMT-based quantitation is the ability to analyze a variety of sample types including primary tissue. However, to realistically monitor ubiquitylation in human cancer tissue and patient-derived cell culture models where generation of protein input of >1 mg/sample is not feasible, a highly-sensitive method that can be successfully employed in the 500 ug - 1 mg range is needed. With this in mind, we have developed a new method for rapid, sensitive and multiplexed ubiquitylation profiling in cells and tissue using anti-K-ε-GG antibodies to enrich ubiquitylated peptides followed by on-antibody TMT labeling. We directly compared on-antibody TMT labeling to conventional in-solution TMT labeling of enriched K-ε-GG peptides and demonstrated that TMT labeling of K-ε-GG peptides while bound to the antibody significantly increases sensitivity, due in part to a significant reduction in the level of TMT contaminant side-products in the final processed sample. We have used this method to quantify over 11,000 ubiquitylated peptides from as little as 500 ug input using only 5.1 h of instrument time, demonstrating that the method is suitable for large-scale studies in primary tissue samples.

Relative to our previously published SILAC-based method for ubiquitylation profiling, the UbiFast method enables multiplexing of >3x more samples, uses 6-12x less peptide input per treatment condition, requires much less wet lab processing time and utilizes 5x less LC-MS measurement time^10^. Additionally, we find that on-antibody TMT labeling increases the relative yield of K-ε-GG peptides relative to SILAC, label free (**Supplemental Figure 2**), and in-solution TMT labeling-based methods. Because of these complexities and inefficiencies in using the SILAC quantification approach for ubiquitylation profiling in cells noted above, studies focused on identifying drug-induced substrates of E3 ubiquitin ligases are being carried out with increasing frequency by proteome profiling only rather than ubiquitin profiling^36,37^. The simplicity and speed of our new on-antibody labeling approach opens the door to deep-scale ubiquitylation profiling to more directly identify the substrates of E3 ligases degraded by the proteasome degradation pathway. Combined with proteome profiling, the UbiFast method should facilitate identification of signaling effects of ubiquitylation as well as substrates of proteasomal degradation. The acetylome and ubiquitylome data sets we provide on the breast cancer xenograft samples are a rich source for the community to begin to probe and elucidate signaling crosstalk between acetylation and ubiquitylation.

A current limitation of the UbiFast method is that the depth of quantification appears to be limited to around 10,000 sites/sample when starting with 0.5-1mg of sample/TMT channel. While this depth-of-coverage is less than what is achieved in SILAC-based experiments, the input amount requirements are lower, the multiplexing capacity is significantly higher, and the method requires no offline fractionation, making it faster and far easier to implement. This is approximately the same depth of coverage achieved using acetylome profiling, but is substantially less than phosphoprofiling that can produce over 35,000 distinct phosphosites/sample in a 10-plex experiment^30^. Extension of our method to higher-plex isobaric labeling methods such as TMTPro (Thermo Fisher) that enables 16-plex analysis should allow further reduction in the amount of sample/channel needed to achieve current depth of analysis. In addition, the quantitative accuracy of the FAIMS-MS2 data generation method we employ is somewhat lower than can be achieved using SPS-MS3, but the lower accuracy is compensated for by the much higher numbers of sites quantified and able to be shown to be differential between states. Future efforts will also be aimed at reducing the amount of labeling reagent required for on-antibody TMT labeling to further reduce the cost of the UbiFast method. Zecha et al have recently presented a protocol for TMT labeling that reduces the quantity of required labeling reagent by reducing reaction volumes^31^.

Beyond applications for ubiquitylation profiling, we hypothesize that on-antibody TMT labeling will be a valuable method to increase the throughput and lower the TMT reagent cost for multiplexed analysis of other PTMs requiring antibody-based enrichment such as lysine acetylation and tyrosine phosphorylation. The UbiFast method is also potentially amenable to serial enrichment analysis workflows^38^ which is especially important when sample amounts are limiting. Finally, we envision that the ease and simplicity of the UbiFast method may be transferable to automated robotic sample-handling platforms to facilitate processing of very large numbers of samples for ubiquitylome analyses^39^.

## Supplemental Figures

**Supplemental Figure 1.**
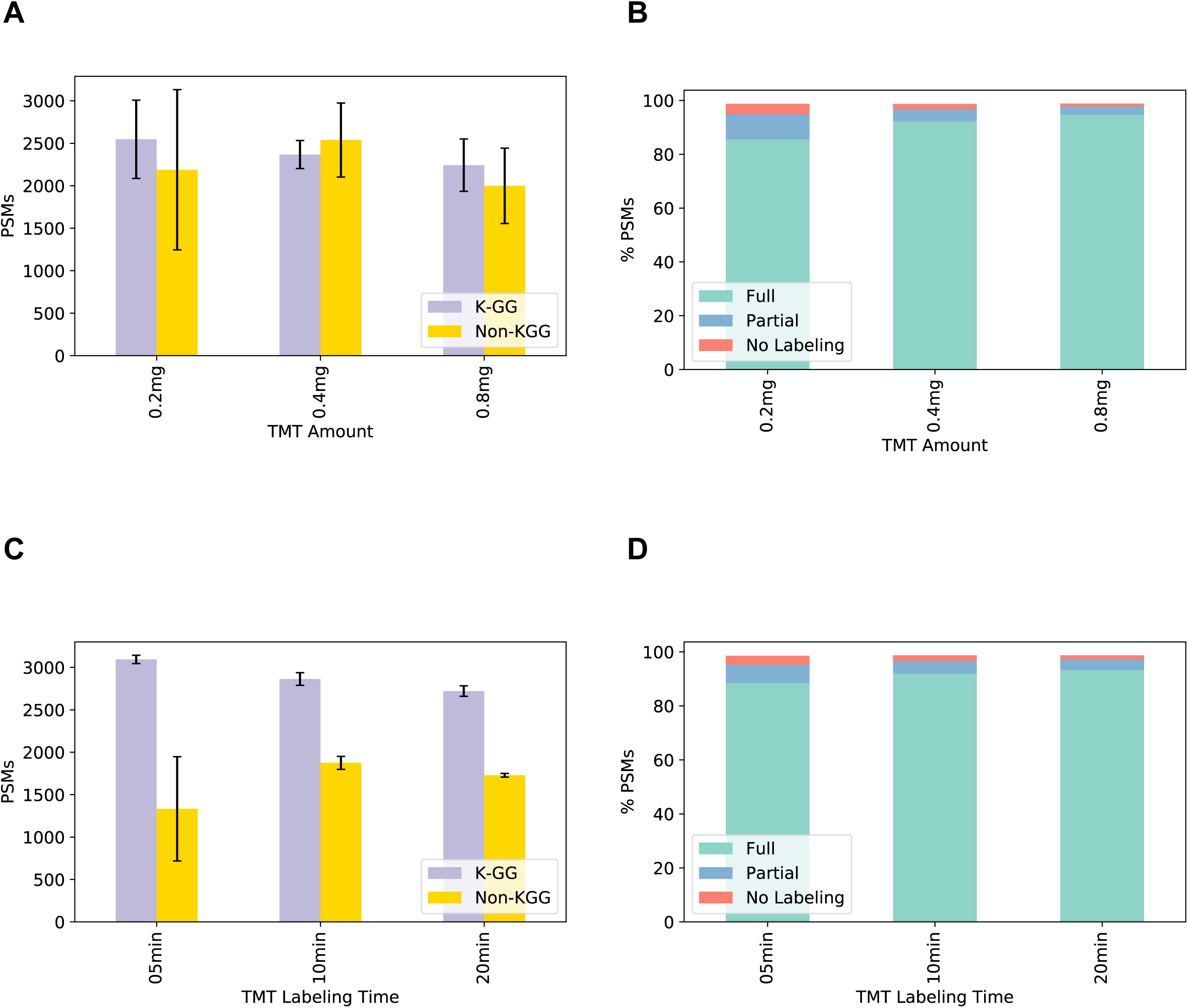
Optimization of on-antibody TMT labeling reagent amount and labeling time. 1 mg Jurkat peptide was processed and labeled in triplicate using on-antibody TMT labeling with a single TMT reagent under the varying conditions shown. The antibody beads were washed to remove non-specifically bound peptides and excess reagent. Bound peptides were then eluted and analyzed by LC-MS/MS using a 110 min gradient. A) Bar plots show the number of PSMs identified using varying amounts of TMT reagent. Error bars indicate standard deviation across three replicates. B) Stacked bar plots show the % of K-ε-GG PSMs fully, partially, or not labeled by TMT for each TMT reagent amount shown. C) Bar plots show the number of distinct K-ε-GG PSMs identified for each TMT labeling time (min). Error bars indicate standard deviation across three replicates, except for the 20 min time point for which duplicate measurements were completed. D) Stacked bar plots show the % of K-ε-GG PSMs fully, partially, or not labeled by TMT for each labeling time (min). K-ε-GG peptides were considered partially labeled if any, but not all, primary amine (i.e. free N-terminus of the peptide and/or side-chain of Lys), other than the K-ε-GG site itself, was labeled by TMT. A peptide was considered to be fully-labeled if all possible primary amines in the peptide (i.e. N-terminus of the peptide and side-chain of Lys), other than the K-ε-GG site itself, were labeled by TMT.

**Supplemental Figure 2.**
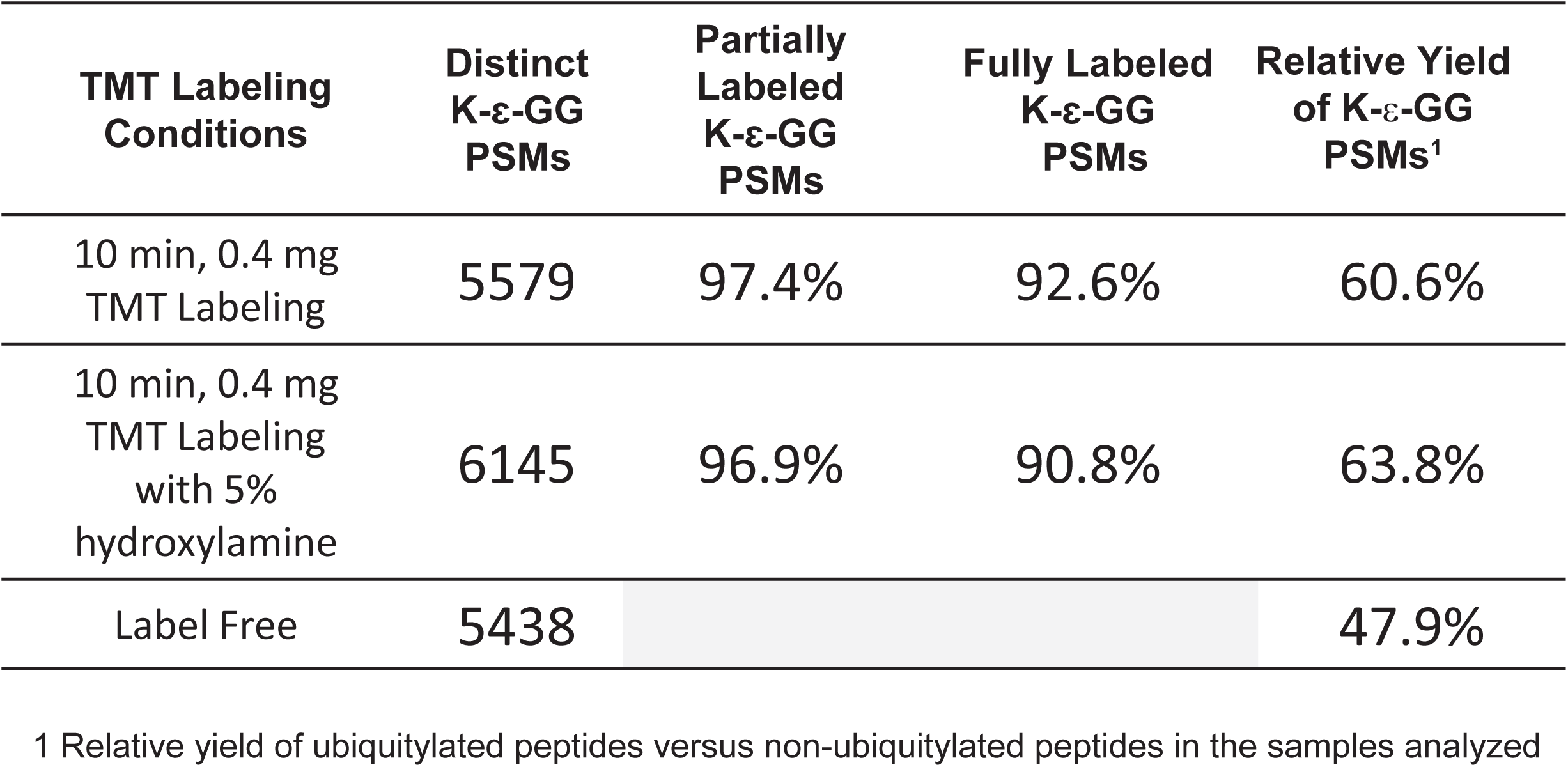
Quenching with hydroxylamine helps prevent TMT cross-labeling using on-antibody labeling. 1 mg Jurkat peptide was processed and labeled in triplicate using on-antibody TMT labeling under the varying conditions shown. The antibody beads were washed to remove non-specifically bound peptides and excess reagent. Bound peptides were then eluted and analyzed by LC-MS/MS using a 110 min gradient. Quenching also increased the number of K-ε-GG peptides identified.

**Supplemental Figure 3.**
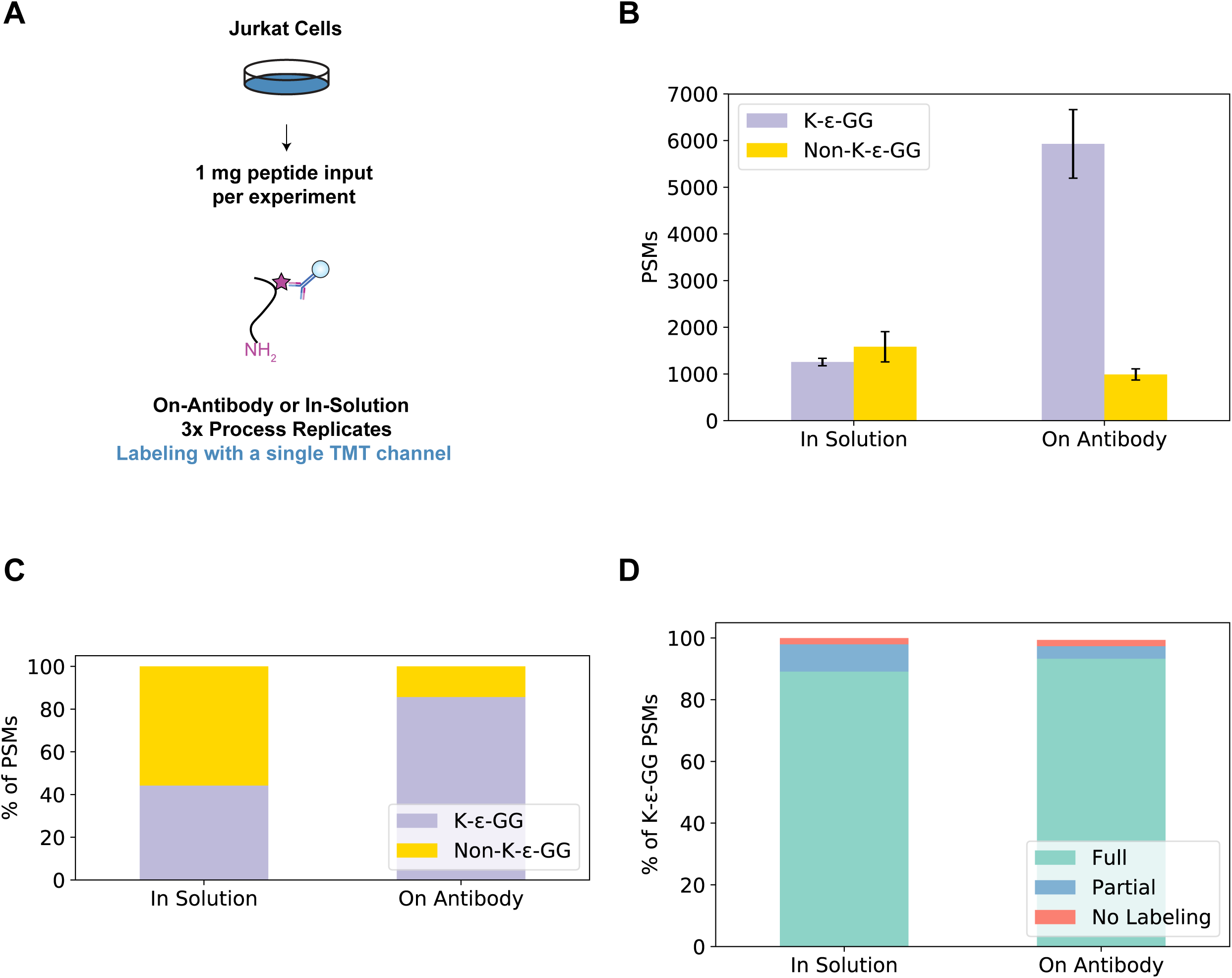
Comparison of numbers of identified K-ε-GG and non-K-ε-GG peptides using on-antibody vs. in solution TMT labeling. A) 1 mg Jurkat peptide was enriched and labeled in triplicate with a single TMT channel using the on-antibody or in-solution TMT labeling method. Each replicate was analyzed by LC-MS/MS using a 110 min gradient. B) Bar plots show the number of K-ε-GG and non-K-ε-GG PSMs for in-solution and on-antibody TMT labeling methods. Error bars indicate standard deviation across three replicates. C) Stacked bar plots show the % of K-ε-GG and non-K-ε-GG PSMs identified by each TMT labeling method. D) Stacked bar plots show the % of K-ε-GG PSMs fully, partially, or not labeled by TMT for each TMT method. K-ε-GG peptides were considered partially labeled if any, but not all, primary amine (i.e. free N-terminus of the peptide and/or side-chain of Lys), other than the K-ε-GG site itself, was labeled by TMT. A peptide was considered to be fully-labeled if all possible primary amines in the peptide (i.e. N-terminus of the peptide and side-chain of Lys), other than the K-ε-GG site itself, were labeled by TMT. The yield of fully TMT labeled peptides was 4% higher using on-antibody labeling versus in-solution labeling.

**Supplemental Figure 4.**
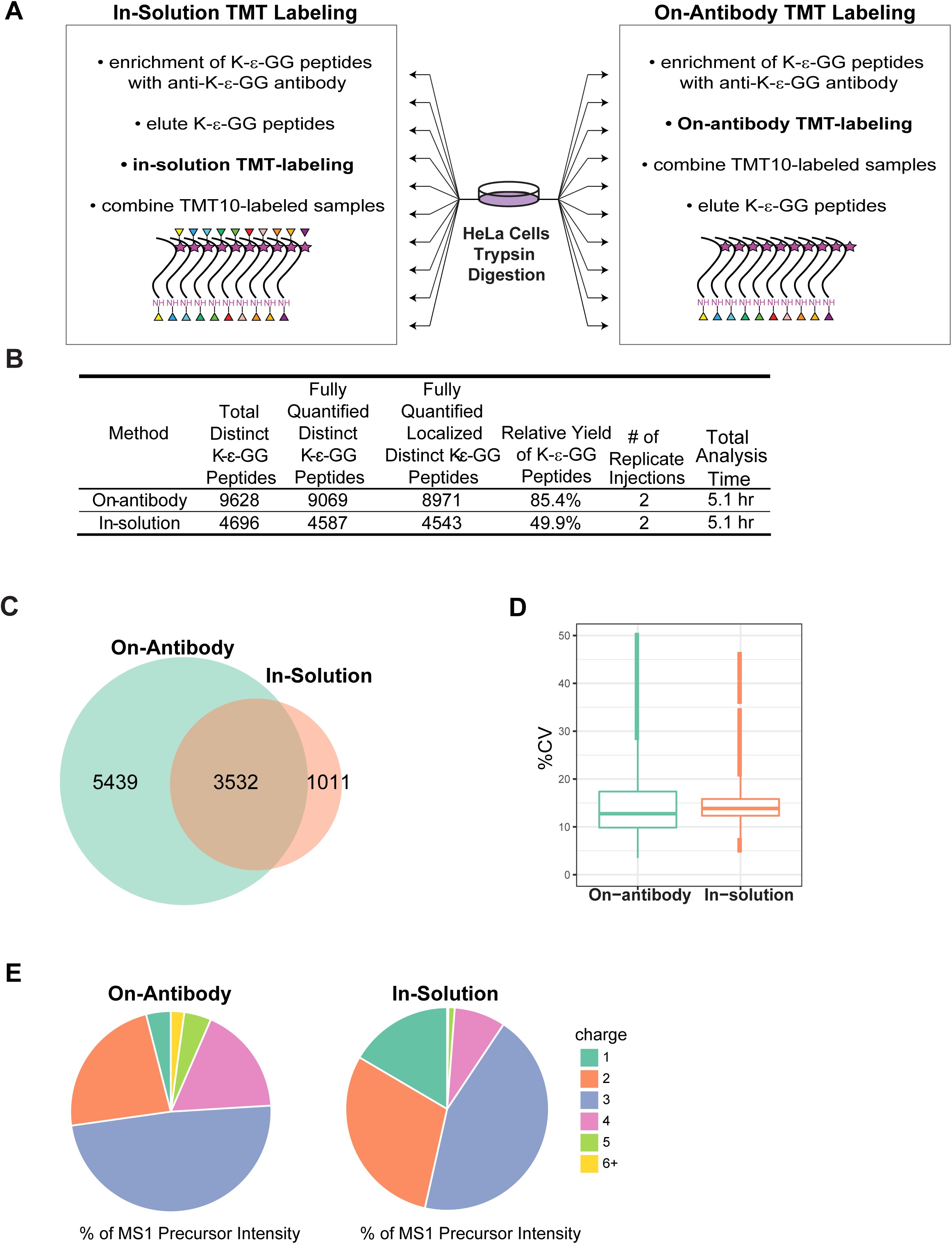
Comparison of on-antibody and in-solution labeling methods A) Experimental design for comparison of TMT10 labeling of ubiquitylated peptides enriched from 1 mg HeLa cells/sample. Labeling was done using the on-antibody method and the in-solution method B) Results of ubiquitylome analyses using the on-antibody and in-solution labeling methods; approximately twice the number of ubiquitylated peptides were obtained using the on-antibody labeling method vs. the in-solution approach. C) Overlap in identified fully quantified, and localized distinct K-ε-GG peptides obtained using the on-antibody and in-solution labeling methods D) Reproducibility (% Coefficients of variation) between HeLa process replicates for each method, and E) % of total precursor intensities separated by charge for each method. Samples were analyzed using a 154 min LC-MS/MS method.

**Supplemental Figure 5.**
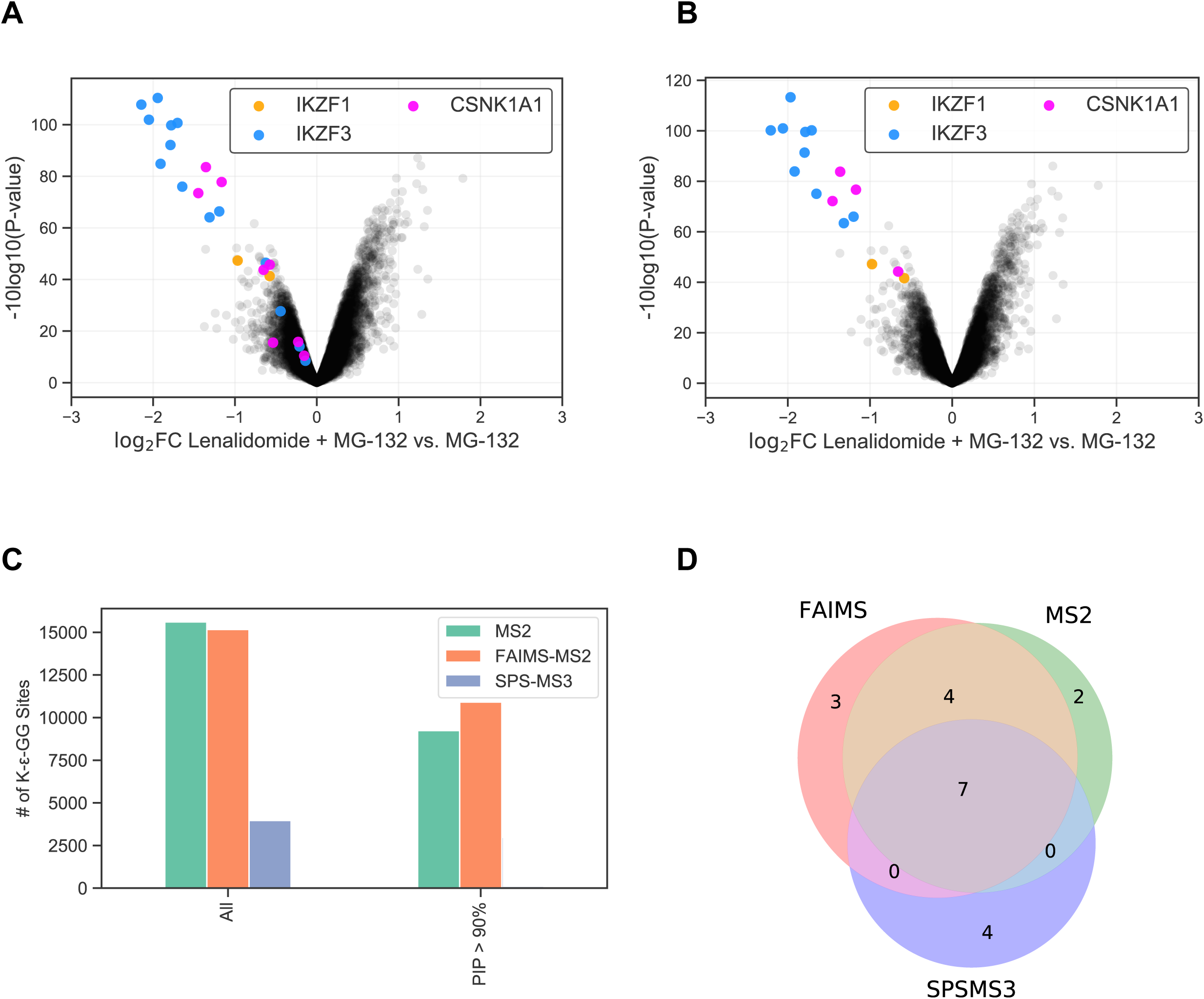
Comparison of MS2, FAIMS-MS2, and SPS-MS3 for the analysis the ubiquitylome of lenalidomide-treated MM1S cells. A) Volcano plot of K-ε-GG sites quantified using MS2; fold change is plotted versus the -log10 of their Padj values following a moderated two-sample t-test. K-ε-GG sites from IKZF1, IKZF3, and CSNK1A1 are colored as indicated in the figure legend B) Volcano plot of K-ε-GG sites quantified using MS2 and filtered for a precursor ion purity (PIP) value of ≥ 90%; fold change is plotted versus the -log10 of their Padj values following a moderated two-sample t-test. C) Bar plots show the number of K-ε-GG sites quantified for each MS method. The second group of bars (right) shows the number of K-ε-GG sites quantified for MS2 and FAIMS-MS2 methods after application of a stringent PIP filter (≥ 90%). D) Venn diagram of the overlap of significant (adj.Pval ≤ 0.001) K-ε-GG sites following a moderated two-sample t-test from IKZF1, IKZF3, and CSNK1A1 across each MS experiment type. For this analysis MS2 and FAIMS-MS2 data were filtered using PIP values ≥ 90%.

**Supplemental Figure 6.**
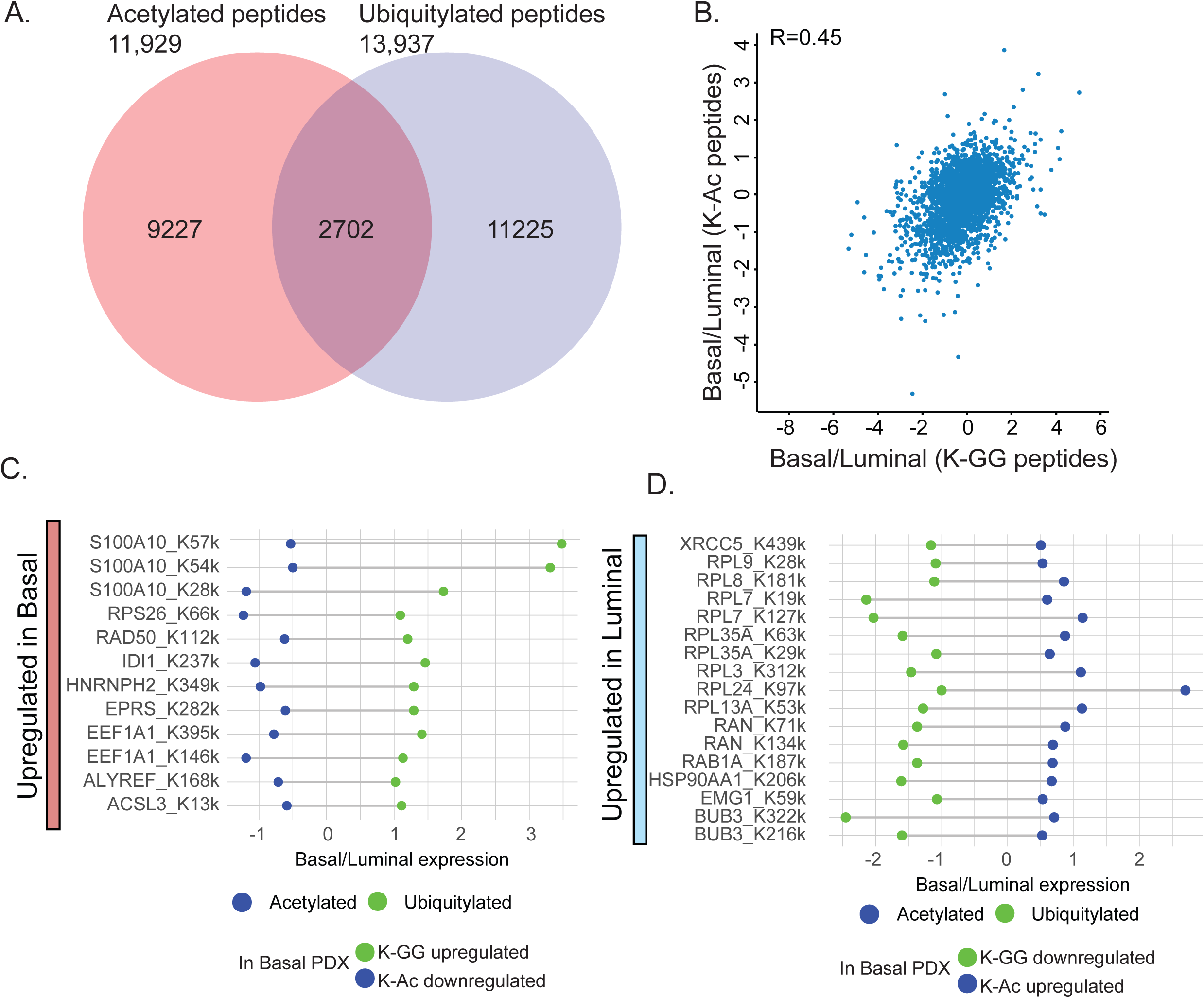
Crosstalk between acetylation and ubiquitylation observed in analyses of Luminal and Basal breast cancer patient-derived xenograft (PDX) samples (see **Figure 3** main text). A) Venn diagram showing the overlap between acetylated and ubiquitylated peptides. B) Scatter plot showing log2 fold-change between Basal and Luminal PDXs, for expression of acetylated and ubiquitylated peptides. C,D) Lolipop plots showing log2 fold-change of acetylated and ubiquitylated peptides between basal and luminal PDXs. The peptides in panel C show specific sites that were upregulated in ubiquitylation and downregulated in acetylation in the basal vs. luminal PDX samples. The peptides in panel D show opposite trend as that of panel C.

## Contributions

N.D.U., D.C.M., P.M. and S.A.C. conceived of the study; N.D.U., D.C.M., SS, T.S., S.F., J.A.G performed experiments; All authors contributed to experimental design, data analysis, and data interpretation.N.D.U., D.C.M., S.S. and S.A.C. generated figures and wrote the manuscript with input from all authors.

## Acknowledgments

This work was supported in part by the following grants: NIH/NCI U24-CA210986, NIH/NCI U01 CA214125 and NIH 1U24DK112340-01 to S.A.C.

